# A dimer peptide ligand of vascular endothelial growth factor slows the progression of human gastric tumors in mouse xenografts

**DOI:** 10.64898/2026.02.18.706549

**Authors:** Xiaoqing Ye, Haofeng Hu, Yilei He, Fei Ye, Jia Jin, Jean-François Gaucher, Lei Wang, Sylvain Broussy

**Affiliations:** Hangzhou Institute for Advanced Study, University of Chinese Academy of Sciences, Hangzhou, China; Université Paris Cité, CNRS, Inserm, CiTCoM, F-75006 Paris, France; College of Life Sciences and Medicine, Zhejiang Sci-Tech University, Hangzhou, China

## Abstract

Gastric cancer is among the most common cancers and represents a major public health problem worldwide. New therapeutic strategies and drugs are needed. Anti-angiogenic agents targeting the Vascular Endothelial Growth Factor (VEGF) are used in combination therapy in the clinic, although their efficacy remains modest. We believe that these large anti-VEGF antibodies could be advantageously replaced by smaller peptides with better tissue penetration. In this study, we evaluate the efficacy of a previously described dimer peptide ligand of VEGF, **D6**, in inhibiting the proliferation of gastric cancer cells and the growth of the corresponding murine xenograft. The activity of the **D6** peptide in these assays was comparable to that of bevacizumab, the positive control antibody, although the peptide required repeated injections at higher molar concentrations. These promising results justify the continued optimization of the peptide dimer, currently under investigation in our laboratory.

## Introduction

Gastric cancer is among the most common cancers, with approximately one million new cases diagnosed worldwide each year. Despite recent advances in treatment, this disease still causes more than 650,000 deaths, and the number of cases in people under 50 is increasing. The incidence of gastric cancer is higher in Asian, South American and Eastern European countries than that in the rest of the world (1). Surgery offers a high chance of survival for operable gastric cancers. In patients with advanced, recurrent or metastatic cancers, systemic chemotherapy is the first line treatment, with possible additional treatments, such as biological agents targeting HER2 and the tight-junction CLDNP18.2 (1).

Vascular Endothelial Growth Factor (VEGF) is the most potent pro-angiogenic cytokine, and inhibition of its signaling pathway is one of the most effective strategies for blocking angiogenesis, a key process in the progression and metastasis of many solid tumors, including gastric adenocarcinoma (2–4). In gastric cancer, the standard second-line treatment includes, in addition to paclitaxel, the anti-VEGF receptor-2 (VEGFR-2) monoclonal antibody ramucirumab, to block angiogenesis. Combinations with immune checkpoint inhibitors are also the subject of numerous studies (1,5). Several tyrosine kinases (TK) inhibitors, including sorafenib, apatinib and regorafenib, targeting the VEGF receptors and other TK receptors with varying degrees of selectivity, have been tested in the clinic as combination therapy, resulting in some cases in beneficial effects, but with significant side-effects (2). For example, apatinib, which selectively targets VEGFR-2, was approved for the treatment of advanced gastric cancer in China in 2014 (6). Recently, the multikinase inhibitor regorafenib has shown improved survival compared to placebo in third line treatment (7). The anti-VEGF antibody bevacizumab has been extensively evaluated in preclinical models and in the clinic in several phase II/III studies for the treatment of gastric cancer (2). Phase II studies have shown that bevacizumab increases the effectiveness of chemotherapy for advanced gastric cancer, although two phase III trials failed to demonstrate any overall survival benefit. The authors note that a benefit may have been observed in non-Asian patients compared to Asian patients (3). Following the results of a phase II/III resectable esophagogastric adenocarcinoma, bevacizumab was not recommended in combination with perioperative chemotherapy, due to the lack of improvement in survival and certain adverse effects (2). These results could be explained by the density of the tumor stroma preventing sufficient perfusion of bevacizumab to the tumor, or by the activation of other angiogenic factors or angiogenic signaling pathways (8). However, a recent retrospective study demonstrated that, in patients with locally advanced gastric cancer, the addition of neoadjuvant bevacizumab to chemotherapy resulted in longer survival, with tolerable adverse events. The authors of the study concluded that its application required further verification (9). Overall, targeting the VEGF pathway to treat gastric cancer is a validated strategy, but it has led to variable results. While monotherapy has proven highly effective in preclinical animal models, combination therapy is required in clinical practice, and results depend on the target (VEGF, extracellular domain of VEGFR or TK domain), drug selectivity and affinity, cancer stage (first, second, or third line), patient history and combination drug. Therefore, it is necessary to develop new molecules targeting the VEGF pathway, exhibiting improved activity and different pharmacokinetic properties. In particular, we believe that tissue penetration properties should be improved. Indeed, the tumor microenvironment is complex, and the uncontrolled proliferation of blood vessels can prevent large biologics, such as antibodies, from reaching the tumor (10).

In this context, our objective is to study new small VEGF ligands capable of blocking its angiogenic activity, with tissue penetration properties different from those of large biologics such as antibodies. To our knowledge, there is no known small molecules capable of directly targeting VEGF and preventing its binding to VEGFRs. Peptides offer a compromise between their size (generally 1 to 5 kDa) and their ability to target such protein-protein interactions. In our ongoing study to optimize VEGF-binding cyclic peptides previously identified by the phage-display technique (11), we shortened their size (12) and improved their affinity by introducing an additional cyclization (13). From these bicyclic monomeric peptides, we synthesized dimeric peptides linked by a polyethylene glycol (PEG) linker capable of targeting the two symmetrical binding sites of the VEGF, which is a homodimeric protein. These peptide dimers bind to VEGF with high affinity, thereby preventing VEGFR activation and blocking VEGF-induced HUVEC proliferation and migration (14). Herein, in order to determine their potential antitumor activity in gastric cancer, the most promising peptide dimer was evaluated on the human gastric cancer cell line SGC-7901 and on a xenografted murine model.

## Results

The **D6** dimer was chosen because it exhibited the highest affinity for VEGF, with K_d,20°C_ = 9 nM and K_d,37°C_ = 26 nM, and the strongest anti-angiogenic activity in HUVEC assays. The new synthesis was adapted from our previously published procedures to provide sufficient quantity to allow *in vivo* experiments at two doses on six mice (14). Monomer **M1** was synthesized from two batches of 0.4 mmol of Rink amide MBHA resin pooled together. The purified monomer was then homodimerized via its Lys 6 side chain with a PEG_25_ linker activated by *N*-hydroxy succinimide groups, to yield 36 mg of **D6** (MW = 5128 g/mol), whose HPLC chromatogram (purity > 98%) and mass spectrometry spectra matched previously published data (Fig 1) (14).

**Figure 1.**
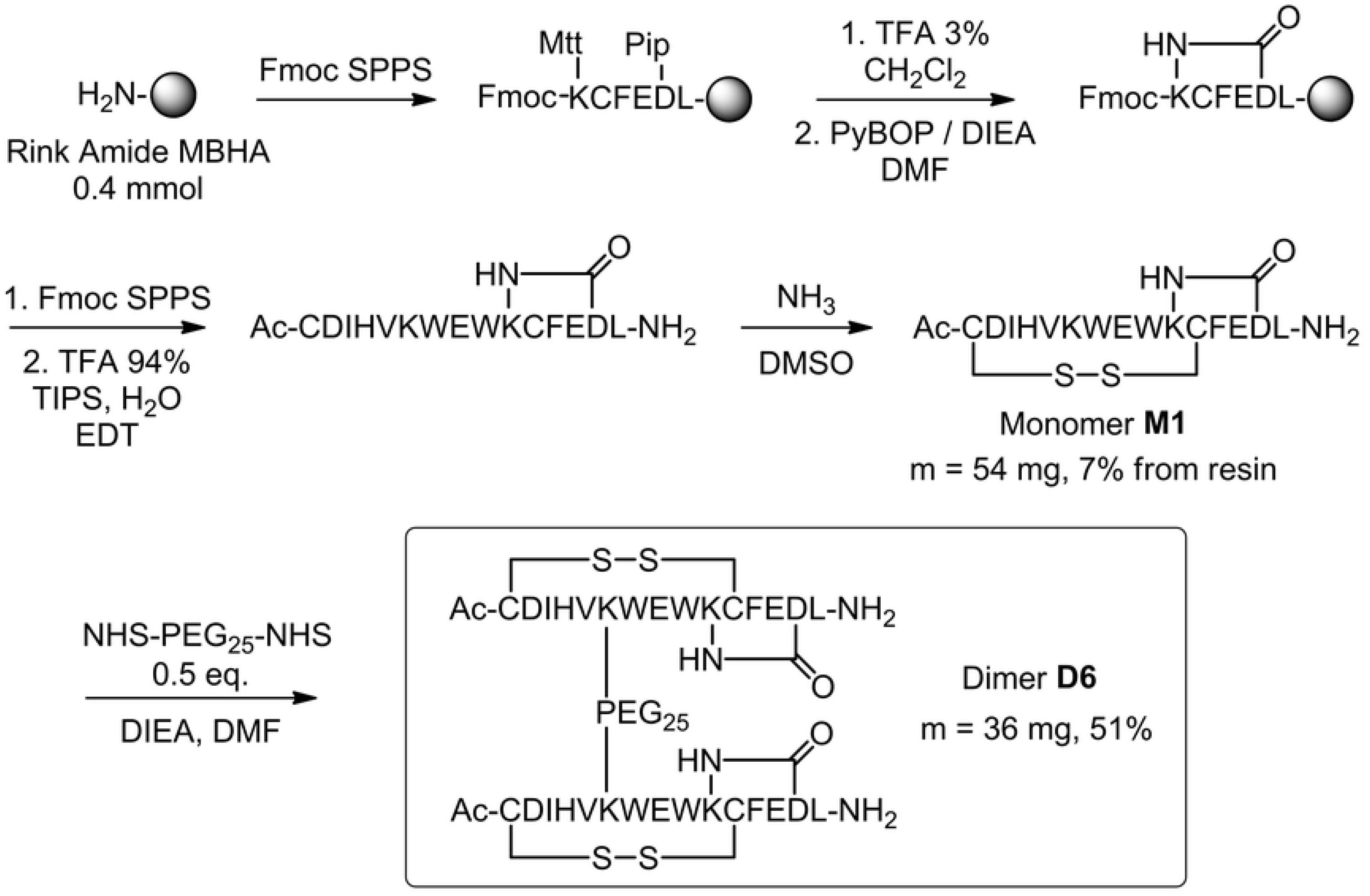
Scale-up (x10) synthesis of the D6 peptide dimer. The **D6** dimer was assayed on the human gastric cancer cell line SGC-7901 (Figure 2). In the groups supplemented with VEGF (50 ng/mL), **D6** induced a dose-dependent inhibition of SGC-7901 cell proliferation at concentrations of 0.4, 2, 10 and 50 μM. At 50 μM, it exhibited antiproliferative activity similar to that of bevacizumab at 6.5 μM, used as positive control. In the groups without VEGF supplementation, no significant effects on cell proliferation were observed, indicating that peptide **D6** did not induce cytotoxicity at concentrations up to 50 μM (Fig 2).

**Figure 2.**
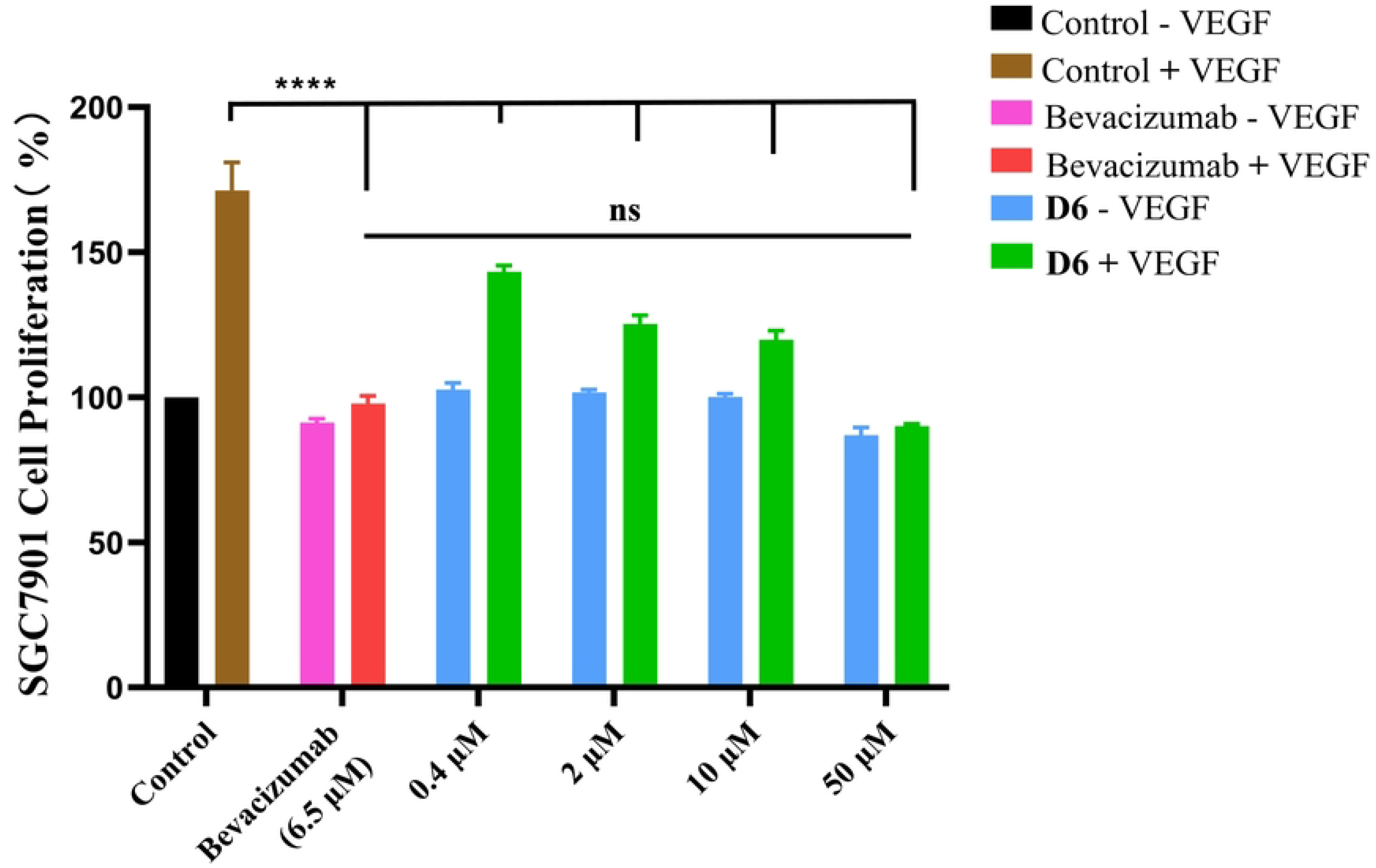
Cellular assays of D6 and bevacizumab on a gastric cancer cell line. Human gastric cancer cells SGC-7901 were treated without (-) or with (+) VEGF, and supplemented with **D6** (at the indicated concentrations), or bevacizumab, or serum-free medium as control groups. Relative cell proliferation (%) was analyzed with GraphPad Prism 8 software. The data are presented as mean ± SD, compared to the control group by one-way analysis of variance (ANOVA). *p < 0.05, **p < 0.01, ***p < 0.001, ****p < 0.0001.

**D6** (5 mg/kg/day and 15 mg/kg/day) and bevacizumab (5 mg/kg, as a single injection, used as a positive control) were administrated by intravenous injection for 2 weeks on BALB/c nude mice bearing a subcutaneous xenograft of SGC-7901 cells. Tumor volume and body weight of mice were measured every 2 days. Peptide **D6** displayed dose-dependent inhibition of tumor growth. Compared to the PBS group, administrated at a dose of 5 mg/kg/day, it reduced tumor volume by 54% and tumor weight by 56%, and administrated at a dose of 15 mg/kg/day, it reduced tumor volume by 68% and tumor weight by 69%. Bevacizumab reduced tumor volume by 75% and tumor weight by 78% with a single administration of 5 mg/kg (Fig 3A, 3B and 3C). During the 2 weeks of the experiment, the inhibitory effect of **D6** administrated at 15 mg/kg/day on tumor volume was comparable to that of bevacizumab administrated at 5 mg/kg once (Fig 3A). No mortality was observed in mice during the 2 weeks of treatment, and monitoring of body weight showed no significant variation compared to the PBS control group, with a slow and steady increase in body weight, as expected (Fig 3D). Therefore, no obvious toxicity of **D6** was observed up to 15 mg/kg/day.

**Figure 3.**
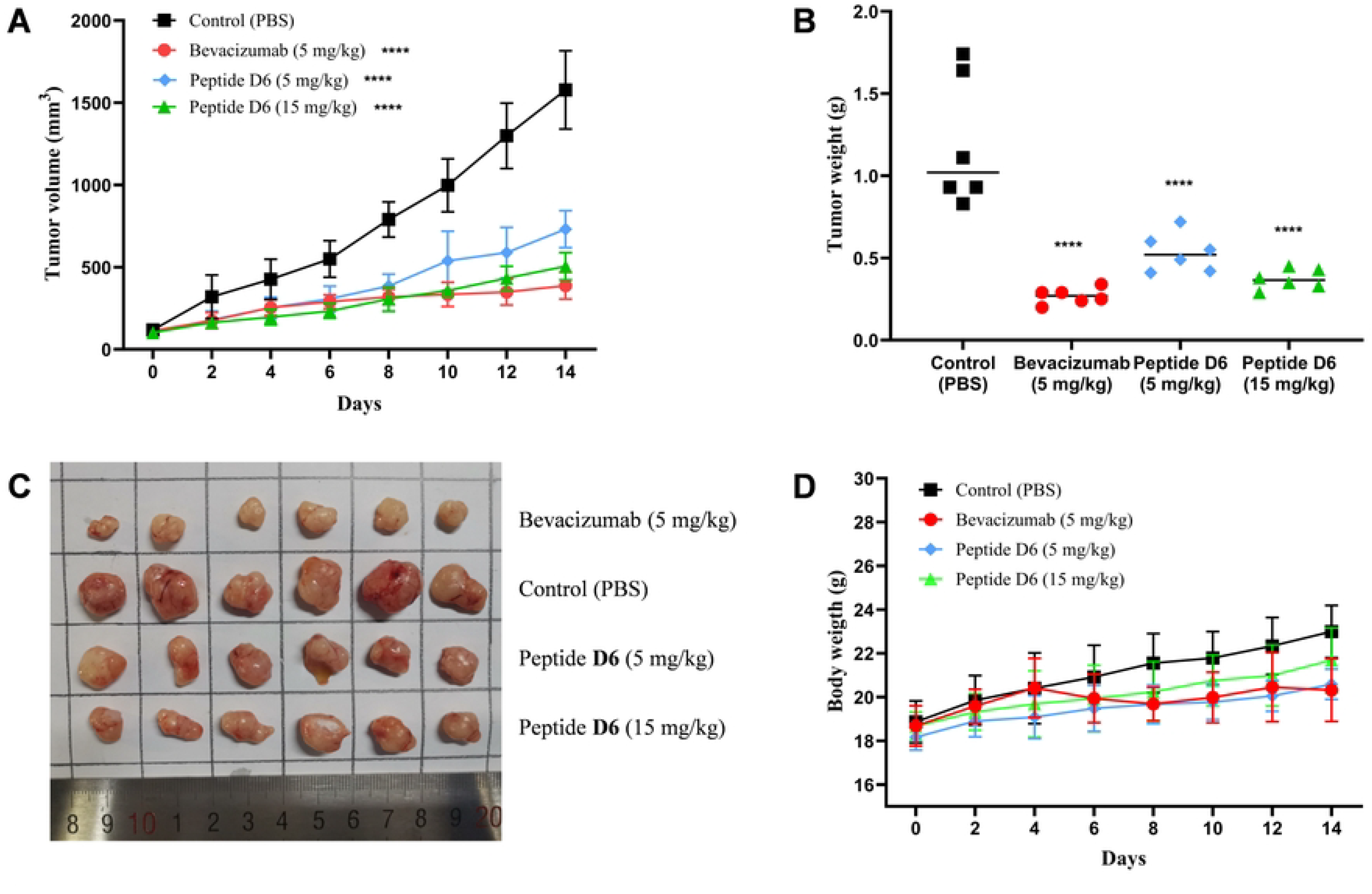
D6 inhibits tumor growth of human gastric cancer cells SGC-7901 on BALB/c nude mice orthotopic transplantation model. (A) Quantification of tumor volume every 2 days. (B) Quantification of tumor weight after 14 days. (C) Images of tumors taken from mice after 14 days of treatment with **D6** (5 mg/kg/day and 15 mg/kg/day), with bevacizumab (5 mg/kg, once) or with PBS (control). (D) Body weight of mice was quantified every 2 days. Data are presented as mean ± SD, compared to the control group by a one-way ANOVA statistical analysis. *p < 0.05, **p < 0.01, ***p < 0.001, ****p < 0.0001.

## Discussion

Angiogenesis is a key process in the growth of solid tumors. Folkman’s initial report suggested that “anti-angiogenesis” treatments should prevent the formation of new blood vessels within the solid tumor, resulting in several therapeutic benefits (15). Since then, anti-angiogenic drugs targeting the main pro-angiogenic growth factor VEGF have been used successfully, often in combination with other drugs, to treat cancers, including gastric cancer (2). There remains a need for new, more potent anti-angiogenic drugs with designed pharmacological properties. In preclinical studies aimed at developing such anti-VEGF drugs, activity assays with on HUVECs and cancers cells requiring this growth factor for their proliferation are standard practice.

We have previously reported a series of VEGF dimer peptide ligands, among which the **D6** dimer showed the best inhibitory activity in HUVEC-based assays (14). Therefore, we tested here its ability to inhibit VEGF-induced proliferation of human SGC-7901 gastric cancer cells and the growth of solid tumor xenografts of the same cell line in nude mice. This cell line was chosen because it is one of the most widely used gastric cell line in preclinical anti-tumor and anti-angiogenic assays, both for cell-based assays and xenografts in mice. It has high mRNA levels of VEGF and VEGFR-2, and requires VEGF for its proliferation (16).

We show here that the binding of **D6** to VEGF was able to effectively suppress the proliferation of SGC-7901 cells in a dose-dependent manner, with inhibitory activity observed from 0.4 µM. The peptide was not toxic to cells up to 50 µM, an important safety feature that has already been demonstrated on HUVEC (14) and on human corneal epithelial cells up to concentrations of 200 µM (HCE-T cells, unpublished results). In mouse xenografts, the **D6** dimer peptide significantly inhibited tumor growth of SGC-7901 cells at a dose of 5 mg/kg/day, and its efficacy at 15 mg/kg/day was comparable to that of bevacizumab at 5 mg/kg. We performed an *in vivo* evaluation of the peptide at doses 5 mg/kg/day and 15 mg/kg/day, as peptides are generally considered to have low *in vivo* stability (17). It is interesting to note that the peptide was administered by intravenous injection, demonstrating its ability to reach the tumor xenograft from the bloodstream. These results can be explained by its bicyclic nature and the presence of the PEG linker, which may improve its *in vivo* stability compared to unmodified linear peptides (18). The promising results obtained with **D6** in mouse xenograft models justify further pharmacokinetic and pharmacodynamic studies.

We have previously described cyclic peptide ligands of the VEGFRs, designed from a VEGF epitope, which have anti-angiogenic activity *in vitro*. Among these peptides, we tested B-cL1 under the same conditions as those used here, *i*.*e*. the SGC-7901 gastric cancer cell line and the corresponding mouse xenograft. The B-cL1 peptide inhibited tumor growth with similar efficacy to bevacizumab at a lower dose of 5 mg/kg/day, which is slightly better than **D6** (19). This result could indicate that targeting the receptor would be a more effective strategy than targeting circulating VEGF in the context of gastric cancer. Bevacizumab (150 kDa) was administered by intravenous injection every 14 days at a dose of 5 mg/kg, the dose usually used in clinical practice (8). Comparing the active concentrations in mol/L of bevacizumab and **D6** (5.1 kDa), it appears that a higher molar concentration of the peptide is required for a comparable therapeutic effect, in agreement with the differences in affinity values measured under the same assay conditions (for the binding of bevacizumab to VEGF, K_d,20°C_ = 0.66 nM) (14). Therefore, new peptides with improved affinities are currently being developed within our research team.

## Materials and methods

### Peptide synthesis

The amino acid coupling, cyclization and dimerization procedures followed the previously published synthesis, with some modifications (14). The scale was increased to two batches of 0.2 mmol of resin, with Rink Amide MBHA resin (substitution 0.78 mmol/g or 0.52 mmol/g) instead of NovaSyn TGR resin (substitution 0.25 mmol/g). The coupling reactions were carried out on an Activo-P14 semi-automatic peptide synthesizer (Activotec) using Nα-Fmoc amino acids (5 equiv relative to the resin) activated with DIC (5 equiv) and OxymaPure (5 equiv) in dimethylformamide (DMF). HPLC analysis was performed using an Uptisphere C18 column (5 µm, 4.6 × 250 mm, Interchim) at a flow rate of 1 mL/min. The purifications were performed by semi-preparative HPLC on a Grace Alltima C18 column (5 µm, 10 × 250 mm, Vydac) with a gradient program from 20 to 100 % B over 40 min and a flow rate of 2 mL/min. The solvents used were water containing 0.1% trifluoroacetic acid (TFA) (solvent A) and a 70% aqueous solution of acetonitrile containing 0.09% TFA (solvent B). The products were detected by UV at 220 and 254 nm.

Briefly, after the synthesis of the first 5 amino acids, the orthogonal protecting groups phenyl isopropyl (Pip) and methyl trityl (Mtt) were removed by treating the peptide resin with 3% TFA in dichloromethane, followed by neutralization with 5% DIEA in DMF. Cyclization was achieved on resin in DMF using 4 equiv of PyBOP and 10 equiv of DIEA relative to the resin, at room temperature for 14 h. Once cyclization was complete, the peptide was elongated to its full length by standard SPPS and acetylated. The peptide was cleaved from the resin with TFA/H_2_O/2,2’-ethylenedioxy diethane thiol (EDDT)/triisopropylsilane (TIPS) (94:2.5:2.5:1.0) for 3 h. The intramolecular disulfide bond was formed by dissolving the crude peptide in dimethyl sulfoxide and adding drops of aqueous ammonia to maintain a basic pH. The reaction was monitored by HPLC. After completion of the cyclization, the crude bicyclic peptide monomer **M1** was purified by semi-preparative HPLC and lyophilized to obtain a white solid. Two batches of monomer peptide were synthesized. In the first batch, 0.2 mmol Rink Amide MBHA resin (0.78 mmol/g) gave 300 mg crude peptide, resulting in 27 mg of pure **M1** (7 % yield). The second batch of 0.2 mmol Rink Amide MBHA resin at 0.52 mmol/g gave 437 mg crude peptide, resulting again in 27 mg of pure **M1** (7 % yield), for a total of 54 mg. The **D6** dimer was synthesized by reacting the peptide monomer **M1**, DIEA (10 equiv relative to peptide), and NHS-PEG_25_-NHS (0.5 equiv relative to peptide) in DMF. The reaction was monitored by HPLC, and the dimer was purified by semi-preparative HPLC, yielding 36 mg of **D6** (51 % yield, purity > 98% by HPLC). The identity of **D6** was confirmed by HRMS with ESI.

### VEGF-induced human gastric cancer SGC-7901 cell proliferation assay

The assay was performed as previously described (19). Briefly, SGC-7901 human gastric cancer cells (2 × 10^3^ cells/well) were seeded onto a 96-well plate in RPMI (Roswell Park Memorial Institute) 1640 medium (Gibco, Life Technologies) containing 2% FBS (Gibco, Life Technologies), and incubated overnight at 37°C, with 5% CO_2_. The medium was then removed and a new serum-free medium, in the presence or absence of 50 ng/mL VEGF-A (R&D Systems) and with different concentrations of **D6** (0.4, 2, 10 and 50 μM), or bevacizumab (6.5 μM, 1 mg/mL, Roche), or control group (serum-free medium) were added and incubated for an additional 48 h (6 wells/concentration/group). Cell proliferation was quantified by Cell Counting Kit-8 (CCK-8, Sigma) assay according to the manufacturer’s instructions. Absorbance was measured at 450 nm using a microplate reader (AMR-100, ALLSHENG, Hangzhou). The experiments were repeated three times.

### *In vivo* antitumor study on a xenografted mouse model

The assay was performed as previously described (19). The evaluation of antitumoral activity was carried out at the Laboratory of Experimental Animal Science, Hangzhou Normal University (Hangzhou, China), maintained under standardized environmental conditions, with protocols approved by the Institutional Animal Care and Use Committee (IACUC) of Hangzhou Normal University. All efforts were made to minimize mice suffering. SGC-7901 human gastric cancer cells (1 × 10^6^ cells/500 μL) from the Cell Bank of the Chinese Academy of Sciences (Shanghai) were injected into the right flank of 8-weeks-old BALB/c female mice. When the tumor size reached 100-300 mm^3^, BALB/c mice were randomly divided into groups (n= 6). **D6** peptide (5 mg/kg/day and 15 mg/kg/day) and PBS (control group) were administrated intravenously for 2 weeks, and bevacizumab (5 mg/kg) was administrated intravenously once, as a positive control. The tumor volume was measured every 2 days with a digital Vernier caliper, using the following formula: v = a^2^ × b × 0.52 (where a is the shortest diameter and b is the longest diameter of the tumor). After 14 days of treatment, the mice were euthanized using CO_2_ followed by cervical dislocation to ensure death, and the solid tumors were separated from the bodies, photographed and weighed (balance OHAUS Adventure).

### Statistical Analysis

The data are expressed as the arithmetic mean ± SEM of at least three different experiments using the GraphPad Prism software version 8.00 (San Diego, United States). The statistical significance of the results was assessed by a one-way ANOVA, with probability values *p < 0.05, **p < 0.01, ***p < 0.001, ****p < 0.0001, being considered significant, “ns” meaning non-significant.

## Acknowledgements

We thank the Laboratory of Experimental Animal Science, Hangzhou Normal University (Hangzhou, China) for animal assay.

## Notes

### Competing Interest Statement

The authors have declared no competing interest.

